# Phylogenetic measures of indel rate variation among the HIV-1 group M subtypes

**DOI:** 10.1101/494344

**Authors:** John Palmer, Art FY Poon

**Affiliations:** Department of Pathology & Laboratory Medicine, Western University, London, Canada; Department of Applied Mathematics, Western University, London, Canada; Department of Microbiology & Immunology, Western University, London, Canada

## Abstract

The transmission and pathogenesis of human immunodeficiency virus type 1 (HIV-1) is disproportionately influenced by evolution in the five variable regions of the virus surface envelope glycoprotein (gp120). Insertions and deletions (indels) are a significant source of evolutionary change in these regions. However, the influx of indels relative to nucleotide substitutions has not yet been quantified through a comparative analysis of HIV-1 sequence data. Here we develop and report results from a phylogenetic method to estimate indel rates for the gp120 variable regions across five major subtypes and two circulating recombinant forms (CRFs) of HIV-1 group M. We processed over 26,000 published HIV-1 gp120 sequences, from which we extracted 6,605 sequences for phylogenetic analysis. In brief, our method employs maximum likelihood to reconstruct phylogenies scaled in time and fits a Poisson model to the observed distribution of indels between closely related pairs of sequences in the tree (cherries). The rate estimates ranged from 3.0 × 10^−5^ to 1.5 × 10^−3^ indels/nt/year and varied significantly among variable regions and subtypes. Indel rates were significantly lower in the region encoding variable loop V3, and also lower for HIV-1 subtype B relative to other subtypes. We also found that variable loops V1, V2 and V4 tended to accumulate significantly longer indels. Further, we observed that the nucleotide composition of indel sequences was significantly distinct from that of the flanking sequence in HIV-1 gp120. Indels affected potential N-linked glycosylation sites substantially more often in V1 and V2 than expected by chance, which is consistent with positive selection on glycosylation patterns within these regions of gp120. These results represent the first comprehensive measures of indel rates in HIV-1 gp120 across multiple subtypes and CRFs, and identifies novel and unexpected patterns for further research in the molecular evolution of HIV-1.

## Introduction

Human immunodeficiency virus type 1 (HIV-1) is a rapidly evolving retrovirus with enormous genetic diversity that is divided into four groups (M, N, O and P). The global HIV-1 pandemic that affects approximately 37 million people as of 2017 [1] is largely caused by group M, which is further partitioned into nine subtypes (A-D, F-H, J, K) that can differ by roughly 30% of their genome sequence [17] and have distinct geographic distributions due to historical founder effects [32]. In addition, there are a large number of circulating recombinant forms (CRFs) that are the result of recombination among two or more HIV-1 subtypes that have subsequently become established in particular regions at high prevalence. The HIV-1 subtypes and CRFs are clinically significant because of variation in pathogenesis, *e.g.*, rates of disease progression, and evolution of drug resistance [6, 16, 37, 38].

In the host cell-derived lipid membrane of every HIV-1 particle, there are numerous virus-encoded envelope glycoprotein complexes composed of three gp41 transmembrane units and three gp120 surface units [35]. The HIV-1 gp120 glycoprotein is a potent surface-exposed antigen that plays a significant role in the recognition and binding of target cell receptors [18]. One reason for the difficulty in immunologically targeting this glycoprotein is the abundance of N-linked glycoslyation sites: sequence motifs that encode the post-translational linkage of glycan groups to asparagine residues [41]. In addition, the HIV-1 gene encoding gp120 has a particularly high rate of evolution, especially within the five hypervariable regions that encode surface-exposed, disordered loop structures. These five variable regions (numbered V1-V5) can tolerate substantially higher amino acid substitution rates than the rest of the HIV-1 genome [19]. Both the extensive glycosylation and rapid substitution rates in HIV-1 gp120 facilitate the escape of the virus from neutralizing antibodies [40].

There are multiple mechanisms by which mutations arise within the HIV-1 genome including nucleotide substitutions, insertions, and deletions [2]. While substitution rates have been extensively characterized in HIV-1 and specifically in the *env* gene [15, 24], less attention has been given to sequence insertions and deletions (indels). The few studies that examine indels in the HIV-1 genome have focused on the location, behaviour, and clinical significance of specifically recurring indels, such as indels in HIV-1 pol associated with drug resistance and indels in *gag* and *vif* associated with disease progression and infectivity [3, 5, 30]. Only a small number of comparative studies have examined indel rates in the HIV-1 env gene encoding gp120 and gp41. Wood et al. [40], for one, found that indels preferentially accumulate in the variable loops of gp120 compared to the remainder of this sequence, while other studies have suggested that variable loop indels correspond with HIV-1 transmission and modulate coreceptor switching [9, 36].

Despite the significant impact of indels within HIV-1 gp120 on virus transmission and adaptation, the overall rates of indel evolution in gp120 have not yet been measured through a comparative analysis. Furthermore, as previous studies on indels in HIV-1 have tended to focus on defined study populations, we have not found any study that has examined indel rates in a large database covering multiple HIV-1 subtypes and geographical regions. Here we present results from a phylogenetic analysis of a publicly available HIV-1 sequence database to estimate the rates of indel evolution in the gp120 variable loops of seven HIV-1 group M subtypes and CRFs (herein referred to collectively as clades). We evaluate the hypothesis that the mean rates of indels significantly vary among the gp120 variable loops and group M clades. Further, we examine the nucleotide composition of indels to assess how this characteristic might be shaped by the virus genome, and quantify the impact of indels on N-linked glycosylation sites in HIV-1 gp120.

## Methods

### Data processing

We queried the Los Alamos National Laboratory (LANL) HIV Sequence Database (http://www.hiv.lanl.gov/) for all sequence records covering HIV-1 env gp120, limiting the records to one sequence per patient. The 26,359 matching sequences were downloaded with predicted subtype, collection year and GenBank accession number. We parsed the resulting FASTA file and removed sequences that lacked subtype or collection year fields, or were shorter than 1400 nt (roughly 90% of full-length HIV-1 gp120), yielding a final data set of 6605 sequences. To extract the interval encoding gp120 from each sequence and partition the result into the variable and conserved regions, we performed pairwise alignments using an implementation of the Altschul-Erickson [4] modification of the Gotoh algorithm in Python (http://github.com/ArtPoon/gotoh2). Each nucleotide sequence was aligned against the HXB2 (Genbank accession number K03455) gp120 reference sequence with match/mismatch scores of +5/–4, gap open/extension penalties of 30 and 10, respectively, and no terminal gap penalty. The aligned query sequence was cut at the boundaries of the aligned HXB2 reference gene to extract the patient-derived subsequence homologous to gp120. Next, we removed any gaps in this result and then aligned the amino acid translation to the gp120 protein reference sequence using an empirical HIV amino acid scoring matrix (25% divergence [23]) with the same gap penalties, except that terminal gaps were penalized at this stage. Finally, we used the aligned query to insert gap character triplets into the preceding nucleotide sequence as ‘in-frame’ codon deletions.

Using the HXB2 reference annotations, we extracted the five variable (V1-V5) and five conserved (C1-C5) regions of gp120. The conserved region sequences were concatenated and exported to separate files for phylogenetic reconstruction. We subsequently determined that our method was not reliably extracting the V5 regions, based on the overabundance of multiple gap characters at the 5’ end of many outputs. To avoid further problems downstream, we implemented a modified extraction method specific to V5. We first extracted nine extra nucleotides beyond the 5’ boundary of the V5 reference to provide conserved sequence coverage outside this hypervariable region. The extended V5 sequence was then translated and aligned to a V5 amino acid reference sequence of matching length as above. Lastly, we used the first non-gap character (a matched amino acid) immediately following the first three conserved residues in the amino acid alignment as the adjusted V5 start position, thereby omitting any gap characters that preceded this first residue.

### Phylogenetic analysis

We used the program MAFFT (version 7.271) with the default settings [14] to generate a multiple sequence alignment (MSA) from the concatenated sequences of conserved regions for each subtype. On manual inspection of the resulting MSAs, we found some alignment columns comprised mostly of gaps caused by rare insertions, so we removed all columns with gap characters in more than 95% of sequences. Next, we reconstructed phylogenies for each subtype-specific MSA by approximate maximum likelihood using FastTree2 (version 2.1.8) compiled with double precision [29]. The resulting trees were manually screened for unusually long terminal branches indicative of problematic sequences, which we removed from the corresponding MSA before reconstructing a revised tree.

Effective estimation of indel rates required that all phylogenetic trees be scaled in time. To rescale the maximum likelihood trees, sequence accession numbers were used to query the GenBank database for more precise collection dates containing month and day fields; otherwise we retained the collection years from the LANL database. The R package *ape* was then used to change each tree into a strictly bifurcating structure and to root the tree using root-to-tip regression [26] based on the associated tip dates. We evaluated the correlation between the time since the inferred root date (x-intercept) and the total branch length (in expected numbers of substitutions) to determine if the data were consistent with a molecular clock [10].

Using the same dates, we employed the least-squares dating (LSD) program [33] to adjust node heights and rescale the tree in time under a relaxed molecular clock model. Dates lacking either month or day fields were specified as bounded intervals. The time-scaled tree outputs from LSD were imported into R to extract the ‘cherries’: pairs of sequences directly descended from a common ancestor with no intervening ancestral nodes. Focusing on sequences in cherries provides phylogenetically independent observations and minimizes the divergence times, thereby reducing the chance of encountering multiple indel events as well as the effect of purifying selection on indels.

### Indel rate estimation

To estimate the rate of indels in the variable loops, homologous variable regions from each pair of sequences in a cherry were compared for length differences. The presence of a length difference was reported as a binomial outcome implying that an indel event had occurred along these branches; this approach does not account for the possibility of multiple indels causing reversion to the same sequence lengths. Additionally, the total branch lengths comprising the cherry was employed as an estimate of divergence time in years. The indel rate was estimated from these data by fitting the following model using maximum likelihood, where the Bernoulli likelihood for the *i*-th cherry is:

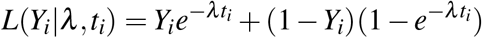

where *Y_i_* = 1 if the sequence lengths differ (implying one or more indels), and is 0 otherwise; *t_i_* is the total branch length, *λ* is the overall indel rate, and *e^−λt_i_^* is the Poisson probability of no indels in the cherry. The total log-likelihood across cherries is thus:

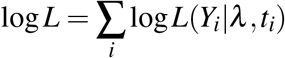

We used the Brent minimization method implemented in the R package ‘bbmle’ to obtain a maximum *λ* for each clade and variable loop combination. A generalized linear model (GLM) with a logit link function was also applied to these data to evaluate statistical associations of the inferred distribution of indels on clades and variable loops; the model incorporated a divergence time term as a rudimentary adjustment for variation in ‘sampling effort’.

### Analysis of indels

For every combination of five variable loops and seven clades, we categorized the inferred lengths of indels into three discrete classes: single-codon (3 nt), double-codon (6 nt), and long (9+ nt). Pearson *χ*^2^ residuals were calculated on these distributions to determine if, and in what direction, these observed proportions significantly deviated from their expected values. To further analyze the composition of indels, we generated pairwise alignments for each cherry with discordant sequence lengths to identify and extract indels. From these pairwise alignments, we calculated the proportions of adenine, thymine, guanine, and cytosine (A,C,G,T) nucleotides in the indel and non-indel regions of the gp120 variable loops. In addition, we recorded the positions and numbers of PNGSs in the five gp120 variable loops by scanning the unaligned amino acid sequences with the regular expression ‘N[ˆP][ST][ˆP]’, where ‘ˆP’ maps to any symbol except P (proline). We then used these data to investigate how commonly indels tended to disrupt PNGSs in the variable loops. By combining PNGS and indel location data, we searched for instances where an indel overlapped with a PNGS and induced to its loss in one of the two sequences of a cherry. To avoid recording instances of partial indel overlap that leave the PNGS intact, we verified the PNGS was disrupted by scanning it again with a regular expression.

## Results

To estimate the rates of indel evolution for different HIV-1 subtypes and circulating recombinant forms (CRFs), we reconstructed phylogenies for each clade using maximum likelihood, and then rooted and rescaled each tree based on the sample collection dates under a molecular clock model. Initially we employed a strict clock model in root-to-tip regressions (Figure 1) to assess whether the data sets contained sufficient signal to estimate rates of evolution (Table 1). Specifically, we confirmed that the lower bounds of the 95% confidence intervals of rate estimates exceeded zero for all clades, which implied a gradual and measurable accumulation of mutations over the sampling time frame. Further, we assessed the model fit with the coefficient of determination (*R*^2^), which was greatest for AE and F1, and lowest for subtype C (Table 1). Next, we employed a more robust least-squares dating method [33] to rescale the trees in time. Table 1 summarizes the substantial differences between the strict clock and least-squares estimates of the times to the most recent common ancestor (tMRCA) for each clade.

**Table 1:** Summary of the evolutionary rate estimates, times to most recent common ancestor (tM-RCAs), and *R*^2^ values generated by applying root-to-tip and least-squares dating models to our seven clade-specific trees. The 95% confidence intervals for the evolutionary rates of both models and for the tMRCAs estimates of the LSD model are enclosed in brackets. Both models are shown to illustrate the differences between fitting strict (root-to-tip) and relaxed clock models to our sequence data.

| Clade | Root-to-tip |  |  | LSD |  |
| --- | --- | --- | --- | --- | --- |
| | Rate $\times 10^{-3}$ | tMRCA | $R^2$ | Rate $\times 10^{-3}$ | tMRCA |
| AE | 2.49 (2.31, 2.67) | 1971.4 | 0.51 | 1.87 (1.85, 2.10) | 1968.6 (1965.5, 1974.6) |
| AG | 2.21 (1.75, 2.67) | 1957.4 | 0.35 | 2.27 (2.10, 2.68) | 1961.9 (1957.5, 1969.0) |
| A1 | 2.69 (2.17, 3.21) | 1932.0 | 0.26 | 2.45 (2.32, 2.66) | 1966.3 (1964.0, 1969.5) |
| B | 2.43 (2.32, 2.54) | 1941.2 | 0.34 | 1.47 (1.46, 1.57) | 1951.7 (1951.2, 1954.8) |
| C | 1.99 (1.75, 2.23) | 1926.9 | 0.13 | 1.80 (1.78, 1.96) | 1939.8 (1937.5, 1946.7) |
| D | 1.90 (1.47, 2.32) | 1944.5 | 0.33 | 1.88 (1.68, 2.11) | 1957.8 (1952.7, 1962.9) |
| F1 | 2.33 (1.85, 2.81) | 1970.4 | 0.57 | 1.67 (1.34, 2.03) | 1956.2 (1943.9, 1965.2) |

**Figure 1:**
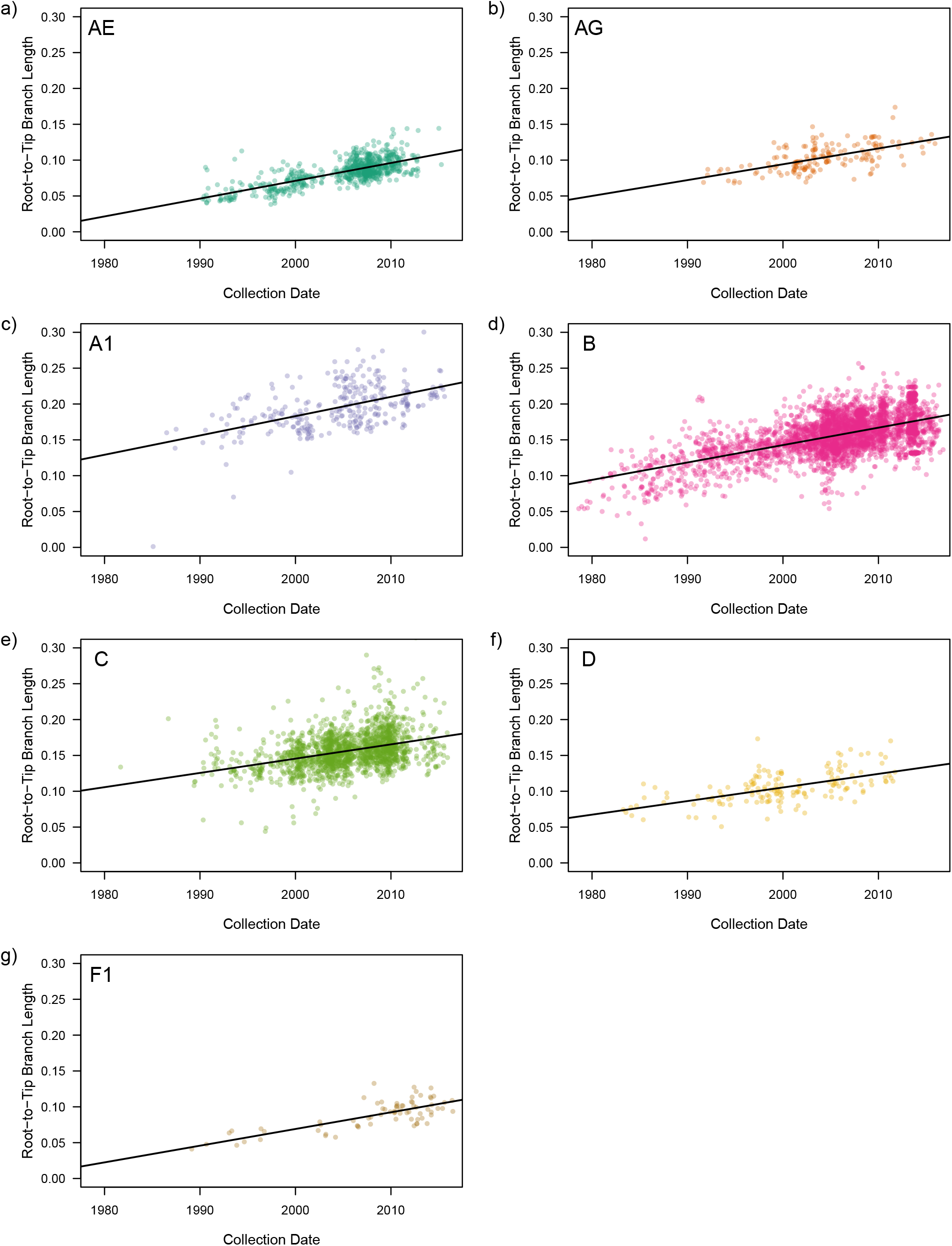
The relationship between the root-to-tip branch length and associated collection dates of sequences within the seven phylogenetic trees generated in this study. The y-axis describes the total branch length from the root to the tips of the tree in units of expected substitutions, while the x-axis describes the collection dates of all recorded sequences. All plots have been adjusted to the same axis scales for comparison. Regions of greater color density indicate the clustering of multiple plotted points. The solid line on each plot describes the linear regression model of the plotted tree tips. (See following page.)

We extracted cherries from these rescaled trees as phylogenetically-independent observations on relatively short time frames and estimated the mean indel rates for each variable loop using a binomial-Poisson model, where the probability of detecting an indel event in a cherry increased exponentially with the divergence time. The indel rate estimates across the five variable loops and seven HIV-1 clades in this study ranged between 3.0 × 10^−5^ to 1.5 × 10^−3^ indels/nt/year (Figure 2). We could not obtain an indel rate estimate for V3 in F1 due due to low sample size for this sub-subtype, such that no cherries had discordant sequence lengths in V3. Similarly, we observed wide confidence intervals for the rate estimates for indels within V1 in AG and F1, and for V5 in F1. The frequency of indels was significantly lower in subtype B than the other clades in our data (binomial GLM, *p* < 2 × 10^−16^; Supplementary (Table S1). In addition, indels were significantly less frequent in V3 irrespective of clade. Estimated interaction effects in the model also indicated that indels were significantly less frequent than expected in V2 within clades B and C.

Under the assumption that differences in sequence lengths of variable loops was caused by a single fixed indel (*i.e.*, no multiple hits), we examined the distribution of indel lengths among variable loops and clades. Cherries with putative indels in the HIV-1 subtype C phylogeny tended to contain significantly longer indels than expected (Figure 3). Conversely, the variable loops V1, V2 and V4 tended to contain longer indels than expected irrespective of clade, whereas V3 and V5 tended to contain shorter indels.

**Figure 2:**
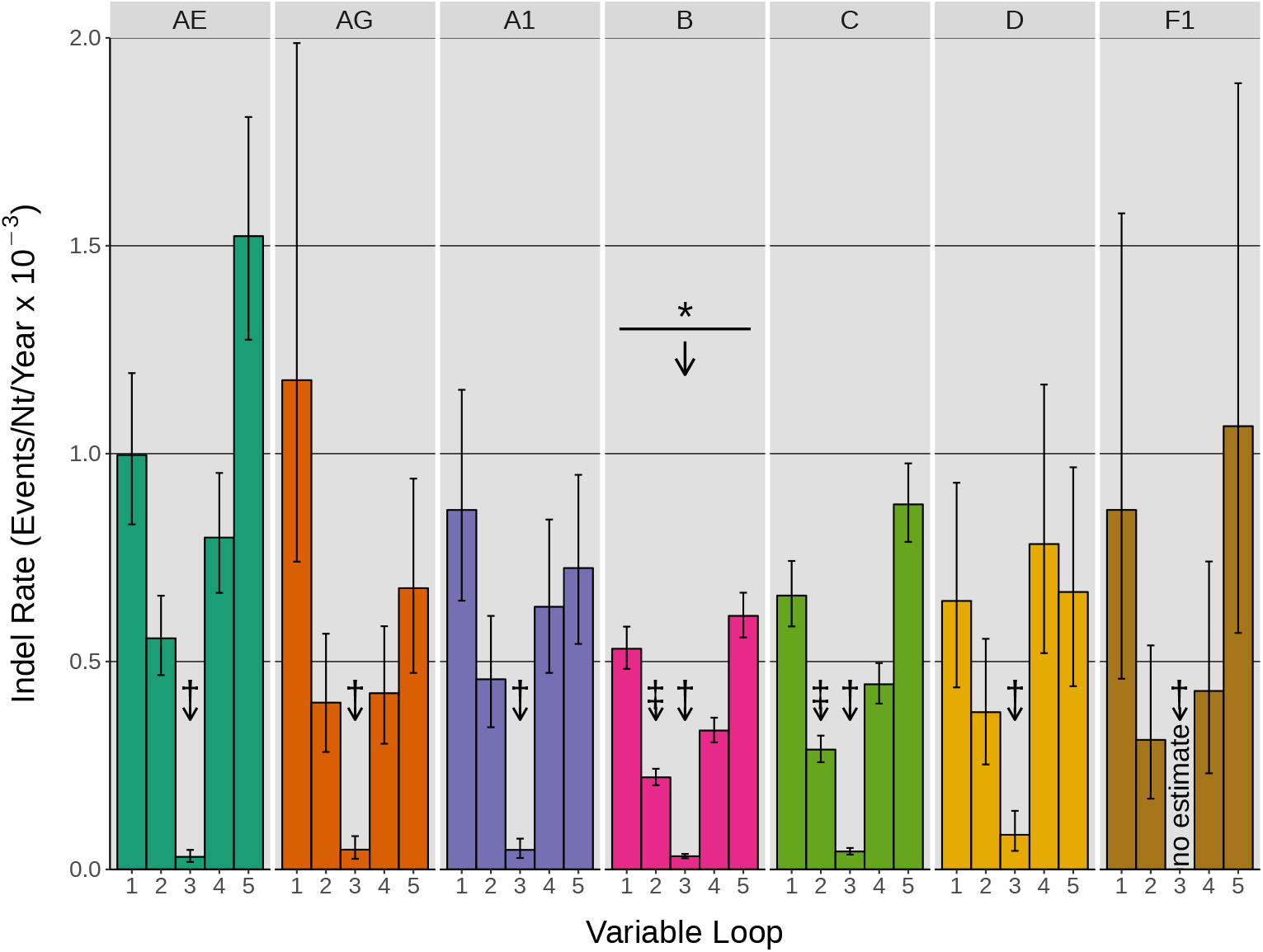
The rate estimates of indel events in the five gp120 variable loops of seven HIV-1 group M clades. Each group of colored bars represents the five variable loop indel rates one of the seven clades. Maximum likelihood estimation was applied to cherry data using a binomial-Poisson model to determine the above indel rates. Error bars represent the 95% confidence intervals within which the indel rates were estimated. Arrows labeled with a * symbol indicate the presence and direction of significant differences among the mean indel rates of group M clades, relative to the CRF AE reference. Arrows labeled by a † symbol denote significant differences among the variable loops irrespective of clade, relative to V1. Arrows labeled by a ‡ symbol denote individual interactions between variable loop and clade which are significantly different than their predicted value. No meaningful rate estimate was provided for V3 of clade F1 because no indels were detected in this data set.

**Figure 3:**
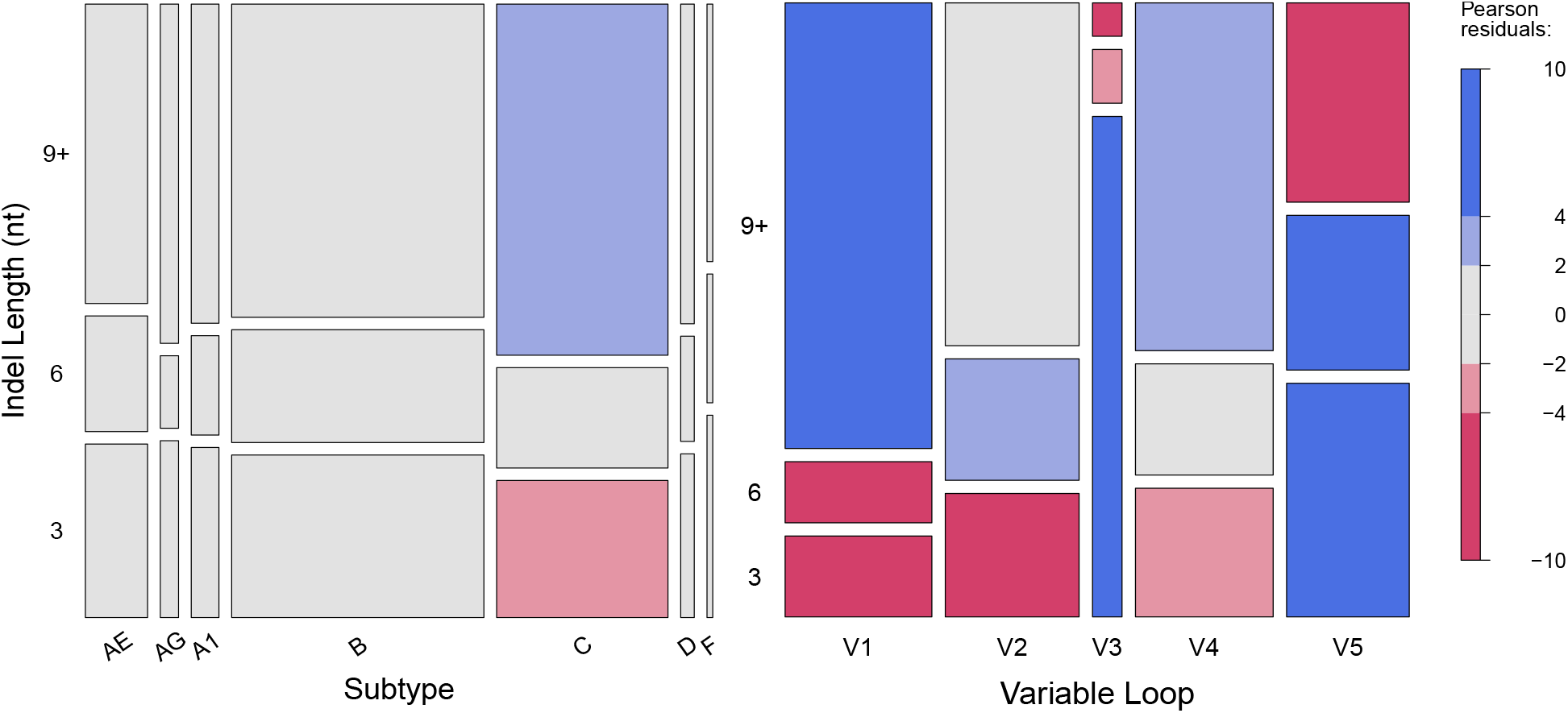
The distribution of indel lengths within the five gp120 variable loops (a) and seven group M clades (b). Indel lengths, measured in nucleotides, were classified into three categories: 3 nt, 6 nt, and 9 nt or longer. Box heights indicate the proportion of indels belonging to the given length category, while box widths indicate the proportion of indels belonging to each variable loop (a) or each clade (b). Pearson *χ*^2^ residuals — quantified measures of the difference between observed and expected values — were calculated for every group on these plots to determine if, and in what direction, these proportions significantly deviated from the *χ*^2^ value. Blue shading represents significantly higher indel counts in a given group than expected based on the Pearson residual, while red shading represents significantly lower counts. Gray boxes did not exceed the Pearson residual threshold of 2 in either direction. Pearson residuals are comparable to the number of standard deviations away from the *χ*^2^ value. As residuals behave similar to standard deviations, values greater than 2, and especially those greater than 4, describe groups whose proportions significantly deviate from the *χ*^2^ value.

Next, we examined the frequencies of nucleotides in indel– and non-indel regions of sequences in cherries with putative indels (Figure 4). Because these frequencies measured for different clades tended to cluster by variable loop, we treated the clades as rudimentary replicates for this comparison (notwithstanding sample variation associated with variable loop V3 and subtype F1, for example). Overall, we observed that indels tended to contain higher proportions of G and lower proportions of T than the corresponding non-indel regions. This pattern was particularly apparent for variable loops V2 and V4.

**Figure 4:**
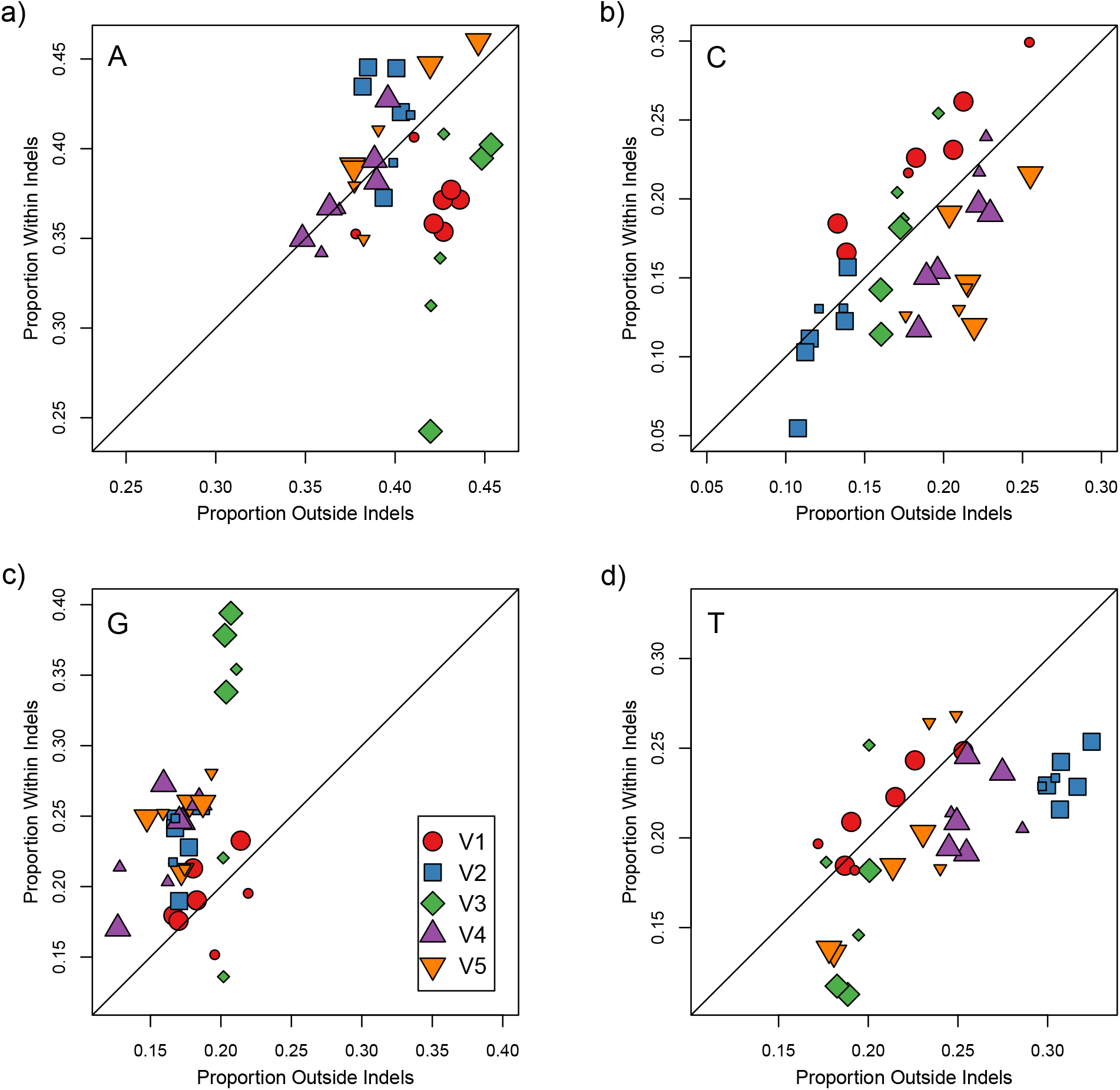
Relations between nucleotide proportions in indel and non-indel sequences for all examined variable loops and subtypes. Plots (a-d) illustrate these relations in adenine, cytosine, guanine, and thymine nucleotides, respectively. Each group denoted by a colored shape represents one of the five variable loops of gp120 and contains seven data points corresponding to each of the examined group M clades. The plotted line with a slope of 1 (y=x) represents the null result in which sequences inside and outside of indels show no difference in their nucleotide proportions. Plotted points that deviate from this line indicate differences between nucleotide proportions found in indels compared to those found outside indels. Larger data points indicate a significant *χ*^2^ test result for that particular data set testing for a difference between indel and non-indel counts.

Figure S1 summarizes the numbers of potential N-linked glycosylation sites (PNGS) for each of the variable loops across the clades in our study. Overall, the loops V1-V5 averaged 2.4, 2.1, 0.9, 4.1 and 1.3 PNGS, respectively. We found significant differences in PNGS counts among clades (Poisson GLM). For variable loop V1, AE contained significantly more PNGS (mean 2.93 PNGS) than the other clades; the next highest count was obtained for subtype C (2.43 [95% C.I. 2.31, 2.57]). Subtypes B and C had significantly higher numbers of PNGS within V3 on average (0.96 [0.87, 1.05] and 0.95 [0.96, 1.05], respectively) than the reference clade AE (mean 0.81). We observed particularly substantial variation in the numbers of PNGS among clades in variable loop V4. For instance, clades AG, A1 and B had significantly higher numbers, and subtype F1 significantly lower, than the reference clade AE (mean 3.72). Finally, we mapped indels to PNGS in the variable loop sequences to determine how frequently indels was associated with the additional or removal of a PNGS (‘disruption’, Figure 5). V1, V2, and V4 contained the highest proportions of indel-induced PNGS disruption among the five variable loops. Again, we observed that estimates for different clades visibly clustered by variable loop. When we adjusted for the relative propotions of the variable loops occupied by PNGS, only V1 and V2 markedly departed from this expectation

**Figure 5:**
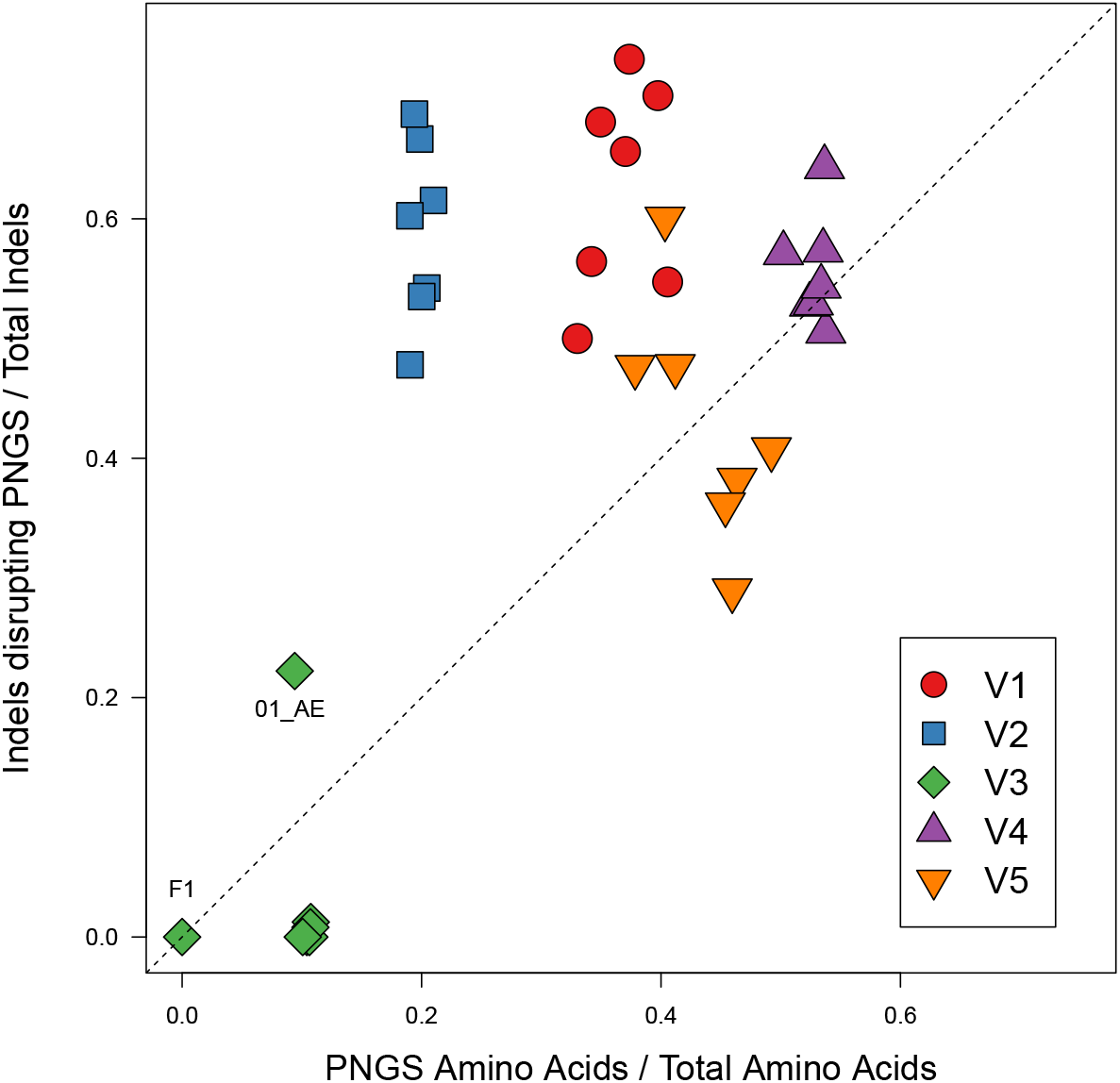
The proportion of indels in a variable region that knocked out at least one PNGS, relative to the PNGS content of the given variable loop. Data sets were represented by a colored shape to denote their variable loop and contained seven points derived from the group M clades. The dotted line of slope 1 provides a rough representation of the general trend expected if PNGSs were under no selection. Individual points that did not cluster with their variable loop were labeled with their clade. Specifically, V3 of subtype F1 did not have any records of indel events.

## Discussion

To our knowledge, these results represent the first comprehensive measurement of indel rates in variable regions of HIV-1 gp120 across major virus subtypes and circulating recombinant forms (CRFs). Surprisingly, one of the only estimates of HIV-1 indel rates we have found dated back to 1995 [22], where Mansky and Temin used an *in vitro* assay of genetic mutations in HIV-1 reverse transcriptase (RT) and reported the observed counts of both nucleotide substitutions and indels. In contrast, our comparative study measures indel rates among different hosts, and as a result will inevitably underestimate these rates due to purifying selection on indels. We sought to reduce the impact of purifying selection by focusing on cherries in the time-scaled phylogenies for each HIV-1 clade. For a given tree, the divergence time between tips in a cherry will be shorter on average than a random selection of two tips, reducing the time that indel polymorphisms were exposed to selection. This emphasis on cherries would have also reduced the probability of multiple indel events accumulating between the lineages in the same variable region. While the comparison of within-host HIV-1 sequences would provide even shorter time scales and thereby more accurate measures of indel rates before selection, some HIV-1 subtypes and CRFs remain underrepresented in publicly available, large and longitudinal same-patient sequence data sets. In addition, results from [40] suggest that purifying selection against indels is relaxed in the variable regions of HIV-1 gp120.

To estimate indel rates, we needed to accurately rescale the HIV-1 phylogenies in chronological time. Estimates of the tMRCA can vary by genomic region, and that estimates from regions within the HIV-1 *gag* and *pol* genes tend to be more recent than regions in *env* [25] irrespective of subtype. Overall, we determined that the diversity of HIV-1 sequences and sample collection dates were sufficient to fit a strict molecular clock model (Table 1). We note that for the purpose of rescaling the trees after this initial assessment with a strict clock, we employed an implementation of a relaxed clock model that allows for rate variation over time. However we also observed that the goodness-of-fit used to assess support for the clock model was the lowest for subtype C. We attribute this poor model fit to both the relatively old age of subtype C [39] and the relative lack of HIV-1 C samples collected prior to 1995 (Figure 1). Estimates of the times to the most recent common ancestor (tMRCA) from the relaxed clock model implemented in the LSD program were generally comparable to previous estimates in the literature for the corresponding HIV-1 clades [11, 39] except for subtype A1, for which we obtained a more recent range of estimates (1964 to 1970). For instance, Tongo et al. [34] recently estimated that (sub-)subtype A1 originated around 1946 - 1957 from an analysis of full-length genome sequence data. We note that because our estimate relies on the ‘point estimate’ of the phylogeny reconstructed by maximum likelihood, the confidence intervals reported for our tMRCA estimates underestimate the true level of uncertainty and fixing the tree may skew the mean estimate. A Bayesian method would more accurately capture this substantial source of uncertainty, but would also be restricted to substantially reduced numbers of sequences due to the complexity of sampling from tree space. Furthermore, Wertheim et al. [39] postulated that estimates of tMRCA among studies may be inconsistent due to the use of nucleotide substitution models that are an inadequate approximation of past molecular evolution.

Our estimates of region– and subtype-specific indel rates ranged from 3.0 × 10^−5^ to 1.5 × 10^−3^ indels/nt/year (Figure 2). As expected, our average estimate (5 × 10^−4^ indels/nt/year) was considerably lower than the rate inferred from Mansky and Temin’s *in vitro* experiments [22] (about 1.5 × 10^−3^ indels/nt/year), where we used parameter estimates from Perelson and Nelson [27] to convert the observed numbers of indel counts to a rate. Since our study compares HIV-1 sequences isolated from different hosts, the indels have been filtered by purifying selection so that only a subset become fixed within the respective hosts. We found that indel rates in subtype B gp120 were significantly lower than the reference clade AE, and generally lower than the other clades in our study. We postulate that this variation of indel rates among subtypes might be associated with differences in the predominant modes of transmission. For instance, more frequent exposure to the mucosal layer, *e.g.*, heterosexual transmission, may episodically select for reduced levels of glycosylation in the HIV-1 envelope to avoid lectin binding [13]. The significantly lower indel rate estimates in V3 irrespective of HIV-1 clade (Figure 2) were consistent with the functional importance of this variable loop. As V3 contributes to HIV entry by binding to the CCR5 or CXCR4 coreceptors, there is substantial purifying selection to conserve its overall structure [12, 20]. This lower tolerance for mutational change, relative to other variable loops of gp120, is consistent with reduced numbers of fixed indels among hosts. The lower indel rates might also be attributed in part to compensatory mutations to preserve structural interactions in V3 [28]. For example, an arginine insertion at position 11 of V3 confers CXCR4 tropism tends to be accompanied by a single amino acid deletion near the C-terminal of V3 [36].

The tendency of HIV-1 subtype C to accumulate longer indels in our analysis is consistent with results previously reported by Derdeyn et al. [9]. By examining HIV-1 heterosexual donor-recipient pairs, Derdeyn et al. [9] first determined that subtype C viruses initially contained shorter V1-V4 sequences upon transmission, which then substantially lengthened by up to 25 amino acids after progressing to chronic late-stage infection. A follow-up study by Chohan et al. [7] provided evidence that this trend was subtype-specific, as it was observed in infections by subtypes A and C, but not subtype B. Similarly, the observed preference for longer indels in V1 and V2 in our data is consistent with the role of these variable loops in facilitating immune evasion. For example, the insertion of five or more amino acids into V1/V2 is associated with reduced sensitivity of HIV-1 gp120 to neutralizing antibodies [8, 31], which is more efficiently achieved by a single long insertion than a series of short insertions. Conversely, the tendency for shorter insertions to accumulate in V3 is consistent with the existence of functional and structural constraints as noted above.

The nucleotide composition of the HIV-1 genome is generally skewed to higher frequencies of A (adenine), in large part due to G-to-A hypermutation induced by host factors [21]. We have not found previous studies that have compared the nucleotide composition of indels to the flanking sequence in the HIV-1 genome. Our analysis indicates that indels in V1 and V3 tend to have substantially higher frequencies of G and lower frequencies of A than the rest of the variable loop (Figure 4). Overall, we observed that indel sequences tended to comprise higher frequencies of G and lower frequencies of T relative to the rest of the variable loop sequence. We note that some frequency estimates had greater sample variation due to limited numbers of indels and sequences in association with V3 and subtypes D and F, for instance. Because the env gene generally contains slightly higher proportions of A (40%) and lower proportions of G (18%) than the rest of the HIV-1 genome (35% and 24%, respectively), we propose that this outcome might reflect the derivation of insertions into env from outlying sequence. Since we have not individually resolved these numerous indels into insertions or deletions through ancestral reconstruction, however, we cannot determine whether this pattern reflects a tendency for sequence insertions to be G-rich, or whether G-rich sequences are specifically targeted for deletion.

The variable regions of HIV-1 gp120 contribute disproportionately to the mean number of potential N-linked glycosylation sites (11 out of 25), which make up the glycan shield of gp120 [42]. Overall, we found that the numbers of PNGS for each variable loop was fairly consistent across clades, with some significant but minor differences in means (Figure S1). Furthermore, we observed that PNGS were more frequently affected by indels in variable loops V1 and V2 irrespective of clade (Figure 5). Put another way, the proportion of indels affecting PNGS in these variable regions was substantially greater than the proportion of the region encoding PNGS. This outcome supports the hypothesis of diversifying selection for the addition or removal of PNGS in V1/V2, where glycosylation plays a major role in mediating immune escape [31] and transmission fitness [9] at different stages in the natural history of HIV-1 infection.

Our estimates of indel rates in the variable regions of HIV-1 gp120 imply that fixed differences in variable loop lengths accumulate between infections on a time scale of about 10-20 years per variable loop with the exception of V3, which accumulates these differences an order of magnitude slower. This time frame is consistent with past observations that HIV-1 gradually ‘raises’ the glycan shield with insertions in V1/V2 several years post-infection [31]. The accumulation and composition of indels among infections is clearly heterogeneous among HIV-1 clades and variable loops. Some of the more exploratory results in this study, *e.g.*, differences in nucleotide frequencies within indels, are particularly novel and suggest new areas for further research in the molecular evolution of HIV-1 to identify the biological or selective determinants of sequence insertions and deletions.

---

**Figure S1:**
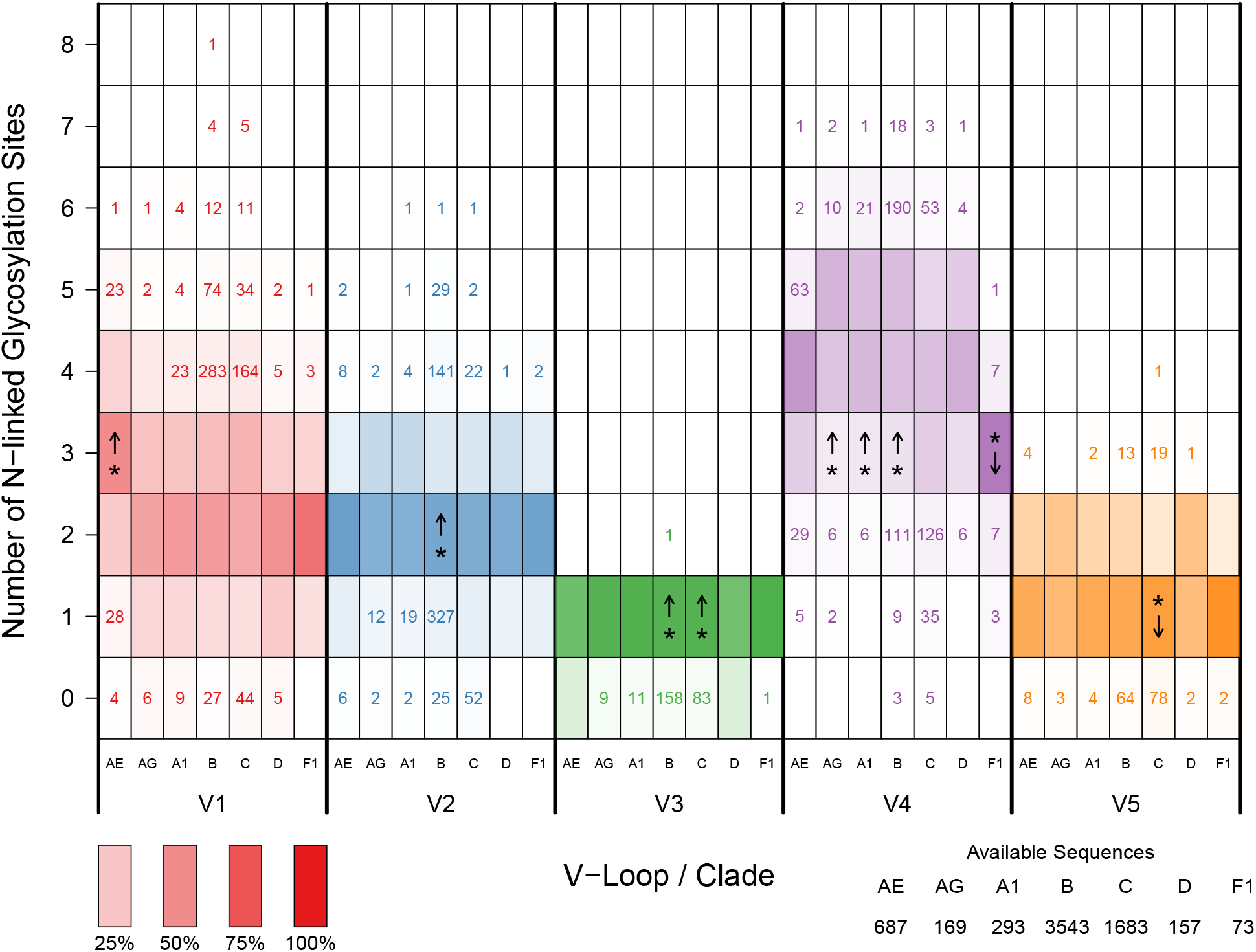
The number of N-linked glycosylation sites in the variable loops of gp120. The color density of each box indicates the frequency of the particular count. Boxes containing a number highlight PNGS counts that are above zero, but contained less than 5% of the distribution and therefore, did not generate a noticeable color. Individual columns within each variable loop section describe the values of the seven evolutionary groups examined in this study. Asterisks (*) and arrows denote subtypes with significantly different distributions and the directionality of these differences within a given variable loop.

**Table S1:** Statistical comparisons generated by applying a generalized linear model to cherry indel analysis. Comparisons made between group M clades and between variable regions were in relation to a reference group (AE, V1). Effects of clade and variable region interactions were compared to predicted mean values to detect significant differences. Groups with asterisk symbols (*) denoted statistically significant differences based on a Bonferroni-corrected threshold suited for multiple comparisons (α = 0.05/n).

|  | Estimate | *P* | Signif. |
| --- | --- | --- | --- |
| Time | 0.001 | $< 10^{-15}$ | * |
| Clade |  |  |  |
| AE | (reference) |  |  |
| AG | 1.946 | 0.058 |  |
| A1 | -0.103 | 0.741 |  |
| B | -1.156 | $1.6 \times 10^{-10}$ | * |
| C | -0.372 | 0.065 |  |
| D | -0.639 | 0.070 |  |
| F1 | -0.158 | 0.780 |  |
| Variable Region |  |  |  |
| V1 | (reference) |  |  |
| V2 | -0.578 | 0.012 |  |
| V3 | -4.448 | $< 10^{-15}$ | * |
| V4 | -0.438 | 0.044 |  |
| V5 | -0.042 | 0.844 |  |
| Interactions |  |  |  |
| AG:V2 | -2.349 | 0.039 |  |
| A1:V2 | -0.457 | 0.313 |  |
| B:V2 | -0.918 | $4.1 \times 10^{-4}$ | * |
| C:V2 | -1.115 | $1.1 \times 10^{-4}$ | * |
| D:V2 | -0.326 | 0.527 |  |
| F1:V2 | -0.860 | 0.285 |  |
| AG:V3 | -2.073 | 0.063 |  |
| A1:V3 | 0.127 | 0.799 |  |
| B:V3 | -0.381 | 0.255 |  |
| C:V3 | -0.109 | 0.752 |  |
| D:V3 | 1.363 | 0.017 |  |
| F1:V3 | -13.565 | 0.926 |  |
| AG:V4 | -2.441 | 0.027 |  |
| A1:V4 | -0.099 | 0.819 |  |
| B:V4 | -0.208 | 0.396 |  |
| C:V4 | -0.215 | 0.428 |  |
| D:V4 | 0.718 | 0.164 |  |
| F1:V4 | -0.577 | 0.439 |  |
| AG:V5 | -2.455 | 0.022 |  |
| A1:V5 | -0.569 | 0.151 |  |
| B:V5 | 0.194 | 0.407 |  |
| C:V5 | 0.246 | 0.345 |  |
| D:V5 | -0.156 | 0.740 |  |
| F1:V5 | -0.188 | 0.797 |  |

